# The basic helix-loop-helix transcription factor TCF4 recruits the Mediator Complex to activate gonadal genes and drive ovarian development

**DOI:** 10.1101/2025.02.28.640455

**Authors:** EV O’Neil, SM Dupont, R Singh, B Capel

**Author notes:** Correspondence: Department of Cell Biology, Duke University School of Medicine.

## Abstract

The bipotential gonad is the precursor organ to both the ovary and testis and develops as part of the embryonic urogenital system. In mice, gonadogenesis initiates around embryonic day 9.5 (E9.5), when coelomic epithelial (CE) cells overlaying the mesonephric ducts proliferate and acquire competence to differentiate into the somatic cell types of the embryonic gonad, the pre-supporting cell (Sertoli cells in the testis and granulosa cells in the ovary) and the interstitial cell lineages. While some transcription factors that drive gonadal cell fate are known, we found that basic helix-loop-helix (bHLH) binding sites are highly represented upstream of granulosa and interstitial genes. We investigated the bHLH Transcription Factor 4 (TCF4), which is expressed in GATA4+ somatic cells in both sexes prior to sex determination, maintained in ovarian pre-supporting cells and interstitial cells of both sexes, but silenced specifically in male pre-supporting cells. In a *Tcf4^STOP/STOP^* mutant mouse model that lacks the TCF4 DNA-binding domain, we found that the marker of Sertoli fate, SOX9, was higher in Sertoli cells, while markers of granulosa and interstitial fate (FOXL2 and NR2F2) were reduced and the supporting to interstitial cell ratio was altered in XX *Tcf4^STOP/STOP^* ovaries. We found that TCF4 binds the Mediator complex and chromatin remodelers to regulate expression of *Jun* and other genes in early somatic cells. Collectively, these results support the working hypothesis that TCF4 regulates a gonadal program that primes the gonad towards a female fate but is specifically silenced in male supporting cells as the testis pathway diverges.

**Significant Findings:**

- bHLH binding sites are more abundant in genes involved in sex-determination than SOX9 or FOXL2 sites, and are 1.5 times more common upstream of granulosa than Sertoli genes.
- TCF4 is a Class I bHLH factor that is expressed in early gonadal progenitors, but silenced in Sertoli cells.
- Loss of the TCF4 binding domain results in higher levels of SOX9, suggesting that TCF4 antagonizes the Sertoli pathway.
- Loss of TCF4 binding domain reduces levels of FOXL2 and NR2F2 and leads to defects in ovary development.
- TCF4 binds with all proteins in the Mediator complex to alter the transcriptional/epigenetic landscape which may explain how it functions on target genes such as Jun.
- Our findings identify *Tcf4* as a potential upstream factor that skews the bipotential supporting cell lineage towards a granulosa cell fate.

## Introduction

Differentiation of the bipotential gonad into either a testis or an ovary is a crucial process with implications for adult reproductive function. In mice, the gonad first forms from epithelial cells on the coelomic surface of the mesonephros around embryonic day 9.5 (E9.5) [1–4]. At this stage, the somatic cells of the XX and XY gonad are bipotential, with virtually identical gene expression and high expression of a common set of gonadal genes including *Wt1, Sf1, Gata4,* and *Lhx9* [5–14].

Sex determination is the process through which the gonad commits to a testis or ovary fate. The testis pathway is initiated by the expression of the Y-chromosome gene *Sry* in a subset of gonadal precursor cells between E10.5 and E12.5 [15, 16]*. Sry* activates the transcription factor *Sox9,* and SOX9 up-regulates a gene expression network that drives differentiation of these cells into Sertoli cells, the testicular supporting cell lineage [17–25]. Differentiation of XX somatic cells into the ovarian granulosa supporting cell fate is associated with expression of an alternate gene regulatory network that includes *Wnt4, Foxl2,* and *Rspo1* [26–30]. In contrast to the male pathway, the upstream factors that drive initial granulosa cell commitment are unclear [31]. This remains a substantial gap in the sex determination field.

An analysis of transcription factor (TF) binding motifs in gonadal cells showed enrichment of basic helix-loop-helix (bHLH) binding sites upstream of granulosa genes (Garcia-Moreno, unpublished data). bHLH proteins are a broad class of TFs that regulate differentiation of cells in many different tissues, including brain and muscle [32–34]. On a molecular level, bHLH TFs typically form homodimers or heterodimers and recognize E-box binding domains (CANNTG) [35, 36]. Of note, the class I bHLH factors, such as MyoD, Twist, and TCF4, are broadly expressed across tissues and are generally associated with transcriptional activation of target genes [37]. In *C. elegans,* the expression of different combinations of class I bHLH TF underly the specification of the sexually dimorphic somatic cell types in the gonad [38]. Whether bHLH factors have a role in sex determination in mammals has not been explored.

In the present study, we investigated whether the class I bHLH factor TCF4 plays a biological role in granulosa cell specification. Our results suggest that TCF4 recruits the Mediator complex to activate gonadal genes in both sexes during early gonadogenesis. While the expression of TCF4 is maintained in interstitial lineages in both XX and XY gonads, TCF4 expression is specifically silenced in XY Sertoli cells, whereas TCF4 is maintained in granulosa cells where it binds with FOXL2 and promotes the female pathway. We suggest a working model in which TCF4 binds in a large activation complex to prime the bipotential pre-supporting cell towards a granulosa cell fate in early gonad development.

## Results

### bHLH binding sites are ubiquitous in up- and down-regulated genes in XY and XX gonads

To determine whether bHLH TFs might play a role in sex determination, we identified genes that were significantly (P < 0.05) up- or down-regulated (≥ 1.15-fold) between E11.5 and E13.5 in the supporting and interstitial cell lineages of XY and XX gonads using a transcriptome dataset from the lab (Supplemental Table 1) [5]. In supporting cells, 43% of the genes that were up-regulated in XX gonads contained HLH binding sites, compared to 29% of the genes that increased in XY supporting cells (Figure 1A). Meanwhile, HLH binding sites were present in 47% of the genes that were down-regulated in XX supporting cells, and in 50% of the genes that decreased in XY supporting cells. Surprisingly, we found that bHLH binding sites were more frequent in the promoter region of the top one hundred most up- or down-regulated genes in both XX and XY supporting cells compared to SOX and FOX TF binding sites (Figure 1A).

**Figure 1.**
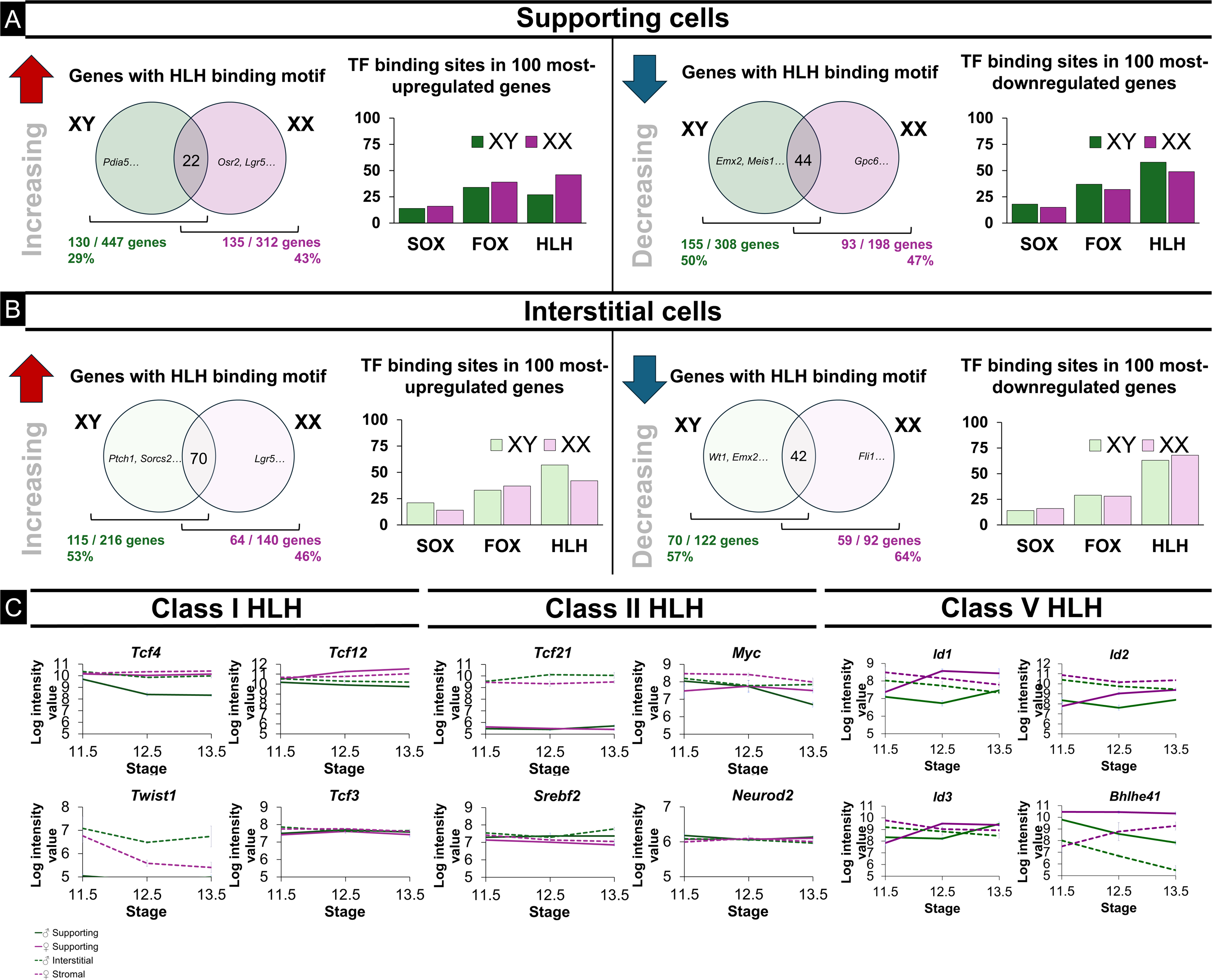
Helix-loop-helix transcription factor binding sites are ubiquitous in dynamically regulated gonadal genes. A) In supporting cells, HLH binding motifs are present in the promoter region of 29% and 43% of genes that increase in XY and XX cells, respectively, and 50% and 47% of genes that decrease in XY and XX cells. Among the top 100 most up- and down-regulated genes in these sets, HLH binding sites are more frequent than SOX or FOX binding sites. B) In interstitial cells, HLH binding sites are present in 53% and 46% of genes that increase in XY and XX cells, respectively. HLH binding sites are enriched in the top 100 most up-regulated genes and occur more frequently than SOX or FOX binding sites. Additionally, HLH binding sites are upstream of 57% and 64% of genes that decrease in XY and XX interstitial cells and are present in nearly 75% of the top 100 most down-regulated interstitial genes. C) Basic helix-loop-helix transcription factors show sex- and cell-type specificity in the gonad (Jameson et al. 2012).

Within the interstitial cell population, HLH binding sites were present in the promoter region of 53% and 46% of up-regulated genes in XY and XX cells. Additionally, HLH binding sites were present upstream of 57% and 64% of down-regulated interstitial genes in the XY and XX gonad, respectively (Figure 1B).

Many HLH transcription factors are expressed within the early gonad (Figure 1C lists representative examples). The Class I bHLH factors *Tcf4* and *Tcf12,* which are associated with gene activation, are highly expressed in both supporting and interstitial cells (Figure 1C). Unlike *Tcf12*, *Tcf4* shows a strong down-regulation in the XY supporting cell lineage. The Class II bHLH TFs are also associated with gene activation but are either specific to the interstitial lineage (*Tcf21*) or generally expressed at lower levels than the Class I bHLH factors. Class V bHLH factors, including *Id1, Id2, Id3,* and *Bhlhe41,* are also highly expressed in the gonad but are associated with gene repression. Given the high expression levels and the activating role of TCF4, along with its significant sexually dimorphic expression pattern, we focused on investigating the role of this bHLH factor in early gonad development.

### The bHLH transcription factor 4, TCF4, is highly expressed in the early gonad and down-regulated in SOX9-expressing cells

As a role for TCF4 in gonad development has not been previously investigated, we sought to first characterize TCF4’s localization in the gonad across gonadogenesis. Using immunostaining (IF), we found that TCF4 was present in the coelomic epithelium and in the broad domain of GATA4+ expressing gonadal cells in XY gonads at E10.75, just prior to sex determination (Figure 2A). At E11.5, expression of TCF4 was excluded from SOX9-expressing cells, but was maintained in XY interstitial cells that were negative for SOX9 (Figure 2B). By E12.5, TCF4 expression became restricted to the interstitial compartment of the testis (Figure 2C), where it co-localized with the interstitial markers, VCAM and NR2F2 (Supplemental Figure 1A). This expression pattern held through E18.5, near the time of birth (Supplemental Figure 1B).

**Figure 2.**
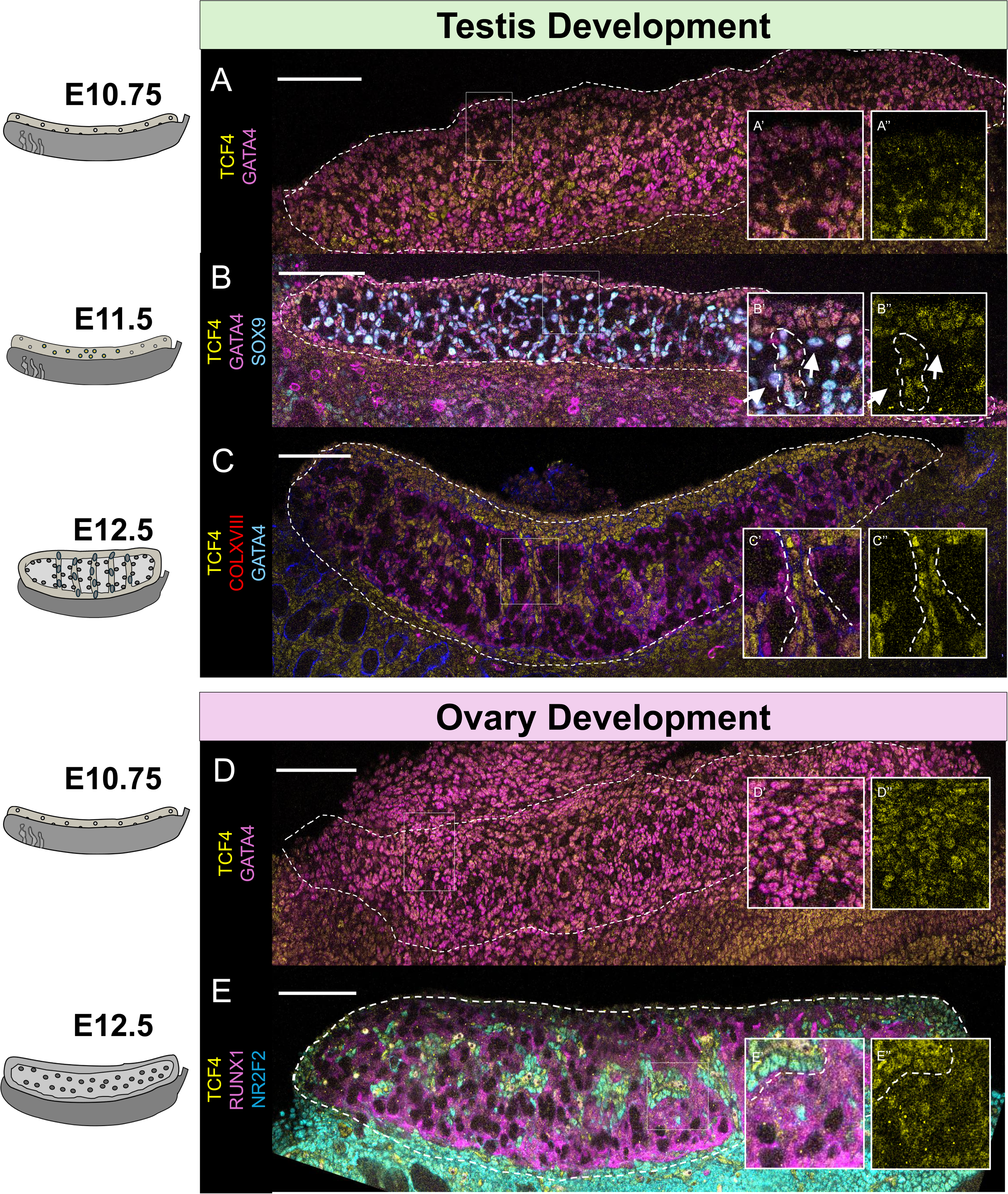
Expression of the HLH transcription factor TCF4 is lost in Sertoli cells but maintained in granulosa and interstitial cell types. A) At E10.75, TCF4 co-localizes with GATA4-expressing gonadal cells in XY gonads. B) At E11.5, TCF4 is lost in SOX9-expressing Sertoli cells in the XY testis, (shown with arrows in B’ and B’’) but is maintained in cells throughout the gonad which are forming sex cords (dotted circle in B’ and B’’). C) In the XY gonad, TCF4 is restricted to the interstitial cell population (shown with dotted lines in C’ and C’’). D) in the XX gonad, TCF4 is co-expressed in GATA4-expressing cells. E) By E12.5, TCF4 is expressed in both RUNX1 expressing granulosa cells and NR2F2+ interstitial cells. Dashed line highlights the boundary between ovarian supporting cells and interstitial cells. Scale bar represents 100 µm. Microscopy images are stitched together from individual panels.

In XX gonads, TCF4 was co-expressed in GATA4+ gonadal cells at E10.75, as in the XY gonads (Figure 2D). However, while TCF4 expression was excluded from Sertoli progenitor cells in the testis, its expression was maintained in both supporting (RUNX1+) and interstitial (NR2F2+) cell types in the developing ovary (Figure 2E). By E18.5, TCF4 was highly enriched in the interstitial lineage, as in the testis, but TCF4 maintained colocalization with granulosa cell markers (Supplemental Figure 1C). Collectively, TCF4 is rapidly down-regulated in Sertoli cell progenitors, but retained in granulosa progenitors and enriched in the interstitial populations of both ovaries and testes.

### Inactivation of the *Tcf4* HLH-DNA binding domain leads to dimorphic effects on the sex determination transcription factors SOX9 and FOXL2

To understand how loss of T*cf4* affects gonad differentiation *in vivo*, we acquired a knock-out-first mutant mouse line carrying a loxP-P2A-GFP-STOP-loxP cassette in intron 17 of the *Tcf4* gene that produces a truncated form of TCF4-GFP lacking the DNA binding domain (Supplemental Figure 2A) [39]. We confirmed the presence of truncated TCF4 by western blot in protein extracts from gonads and limbs of *Tcf4^STOP/+^* and *Tcf4^+/+^* fetuses (Supplemental Figure 2B). Offspring homozygous for this disruption of *Tcf4* die a few days post-birth, which restricted our analysis of mutants to fetal stages. Mutant *Tcf4^STOP/+^* studs were crossed with *Tcf4^STOP/+^* dams, and *Tcf4^+/+^* and *Tcf4^STOP/STOP^* littermates were collected for downstream analysis.

First, we performed immunostaining to determine if general gonad development was disrupted by *Tcf4* inactivation. XY wildtype *Tcf4^+/+^* (n = 3) and mutant *Tcf4^STOP/STOP^* (n = 6) littermates showed relatively normal development of the testis, with organization of germ cells into sex cords surrounded by SOX9+ Sertoli cells (Figure 3A). Additionally, HSD3B1+ interstitial cells were present by E15.5, suggesting that interstitial cell types differentiated normally. In XX *Tcf4^+/+^* (n = 5) and *Tcf4^STOP/STOP^* (n = 5) fetuses, ovary development also progressed relatively normally, with the appearance and organization of FOXL2+ lined ovigerous cords (Figure 3B). However, some *Tcf4^STOP/STOP^* mutant ovaries had pockets of FOXL2+ granulosa cells lacking germ cells, suggesting that there could be disruption to ovarian organization in these mutants (Supplemental Figure 3A).

**Figure 3.**
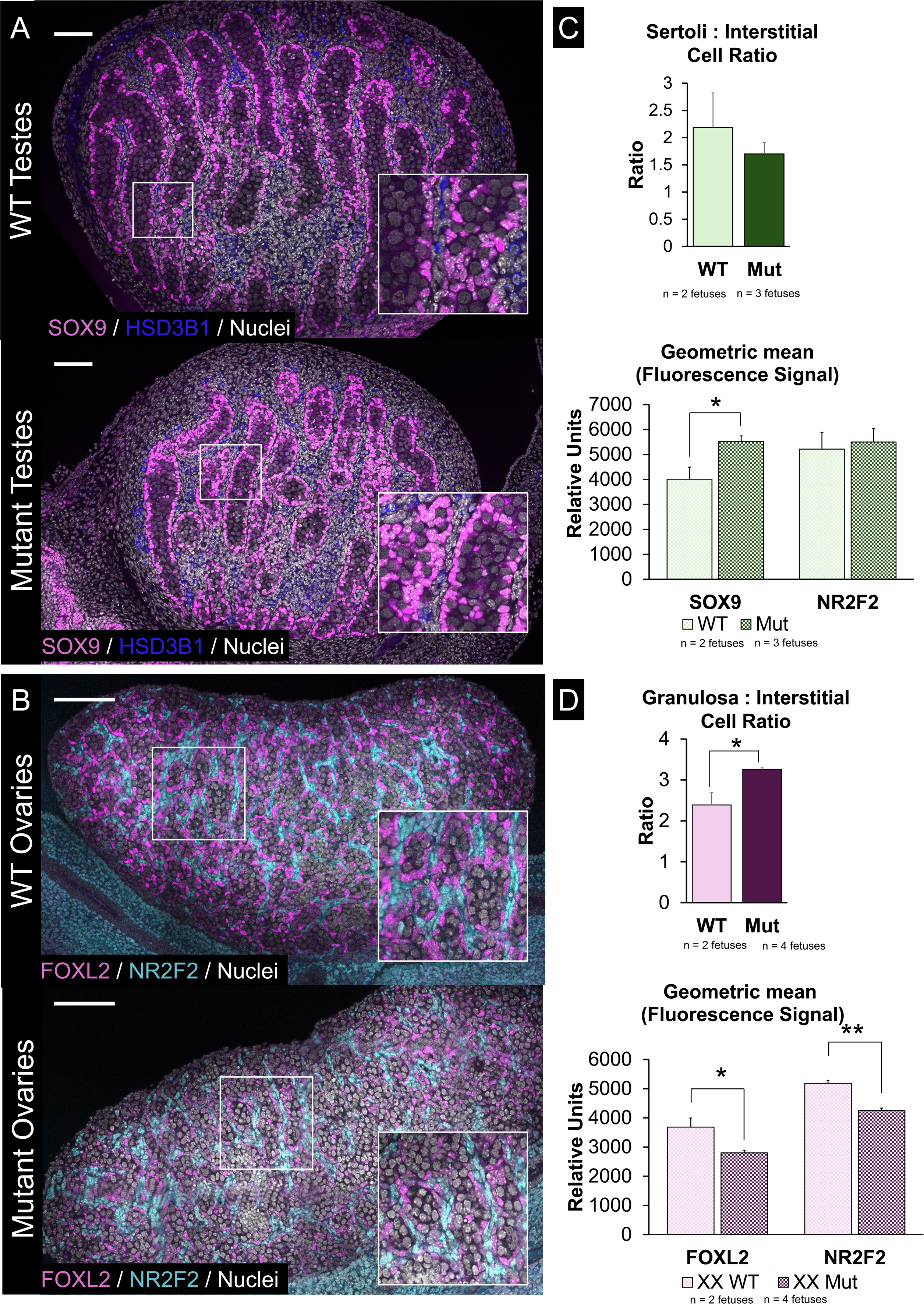
TCF4 does not disrupt gonad differentiation but does affect expression of master regulators of sex determination and cell type ratios in the ovary. A) Immunofluorescence imaging of wildtype *Tcf4^+/+^* and mutant *Tcf4^STOP/STOP^* XY gonads. Images were taken using identical microscope settings. Representative images shown. B) Immunofluorescence imaging of wildtype *Tcf4^+/+^* and mutant *Tcf4^STOP/STOP^* XX gonads. Images were taken using identical microscope settings. Representative images shown. Microscopy images are stitched together from individual panels. C) Gonads were digested and run on a cell analyzer. While ratios of supporting: interstitial cell types did not change, SOX9 was more abundant (P < 0.05) on a per-cell bases in *Tcf4^STOP/STOP^* XY gonads (n=3 gonad pairs) relative to *Tcf4^+/+^* gonads (n=2 gonad pairs). Meanwhile, NR2F2 was not different between samples. D) Mutant ovaries had a higher supporting: interstitial cell ratio (P < 0.05). Additionally, FOXL2 and NR2F2 were less abundant (P < 0.05) on a per-cell bases in *Tcf4^STOP/STOP^* XX gonads (n = 4 gonad pairs) relative to *Tcf4^+/+^* gonads (n = 2 gonad pairs).

Since bHLH factors specify gonadal cell types and regulate sex determination in invertebrate species [38], we used FACS analysis to test whether loss of TCF4 alters the proportions of supporting and interstitial cells in E15.5 testes and ovaries. Gonad pairs from single fetuses were dissociated, fixed, and stained for markers for supporting cells (SOX9 in XY or FOXL2 in XX) or interstitial cells (NR2F2).

Use of FACS analysis allowed us to measure cell numbers and protein abundance using fluorescent readings on a single-cell level. The proportions, ratios, and geometric fluorescence values of *Tcf4^+/+^* and *Tcf4^STOP/STOP^* gonads are listed in Supplemental Table 2.

XY *Tcf4^STOP/STOP^* mutant gonads (n = 3 XY gonad pairs over 3 litters) had similar proportions of supporting and interstitial cells relative to wildtype *Tcf4^+/+^* XY gonads (n = 2 XY over 2 litters). However, there was higher expression of SOX9 protein on a per-cell basis in Sertoli cells of *Tcf4^STOP/STOP^* mutants (n = 10,000 cells per gonad pair, 3 gonad pairs over 3 litters) compared to *Tcf4^+/+^* littermates (n = 10,000 cells per gonad pair, 2 gonad pairs over 2 litters) (P < 0.05). There was no difference in levels of the interstitial transcription factor, NR2F2, between wildtype and mutant gonads (Figure 3C).

Closer analysis of cell types in the ovary on the BD LSRFortessa™ Analyzer showed that mutant *Tcf4^STOP/STOP^* ovaries (n = 10,000 cells per gonad pair, 4 gonad pairs over 3 litters) had a greater supporting to interstitial cell ratio relative to XX *Tcf4^+/+^* wildtype littermates (n = 10,000 cells per gonad pair, 2 gonad pairs over 2 litters). Further, we observed decreases in the average level of both FOXL2 and NR2F2 (P < 0.05) protein on a per-cell basis relative to wildtype *Tcf4^+/+^* ovaries (Figure 3D).

### Across both sexes, TCF4 recruits the Mediator Complex and other core transcriptional machinery to regulate gene expression

Given the effects of TCF4 inactivation on ovarian development, we sought to further investigate TCF4’s function at E12.5 by characterizing binding partners and gene targets around the time of pre-supporting cell commitment to the granulosa cell fate. Using the Nr5a1-GFP line, which labels both supporting and interstitial cells at E12.5 [5, 40], we used FACS to isolate cells, immunoprecipitated TCF4, and performed mass spectrometry (MS) (Figure 4A) (Supplemental Table 3A, B, and C). Binding partners of TCF4 were defined as proteins that showed at least a 1.5-fold difference between the TCF4 IP and IgG IP in each sex. In total, 2,215 proteins co-precipitated with TCF4 in dissociated XX gonadal cells, and 3,130 proteins immunoprecipitated in dissociated XY cells as defined by a significant enrichment (P = 0.05, rounded to two decimal places) relative to the IgG control IP. Of these, 1,774 proteins co-precipitated with TCF4 in both XX and XY gonads (Figure 4B). Notably, TCF4 itself was enriched in the TCF4 IP of both XX (P = 0.05) and XY cells (P = 0.04). The majority of TCF4 binding partners common to both XX and XY gonads were not differentially abundant by sex (96%) and the HLH factor TCF21, an interstitial-specific bHLH TF [41], immunoprecipitated with TCF4 in both XX and XY cells. Shared TCF4 binding partners in both XX and XY cells were enriched in factors involved in positive regulation of transcription (Figure 4C). Many key regulators of basal transcriptional processes co-precipitated with TCF4, including all 24 of the Mediator complex proteins, as well as several classes of epigenetic regulators, co-activators, transcription factors, and the RNA polymerase holoenzyme (Figure 4D). We independently confirmed the presence of one core Mediator protein, Mediator 8, via western blot analysis of a TCF4 pulldown (Figure 4E). While most proteins precipitated with TCF4 in both XY and XX cells, there were also sex-specific TCF4 binders (Figure 4F). Most notably, the core transcription factor, FOXL2, which has well established roles as a crucial regulator of granulosa cell identity, co-precipitated with TCF4 in XX cells.

**Figure 4.**
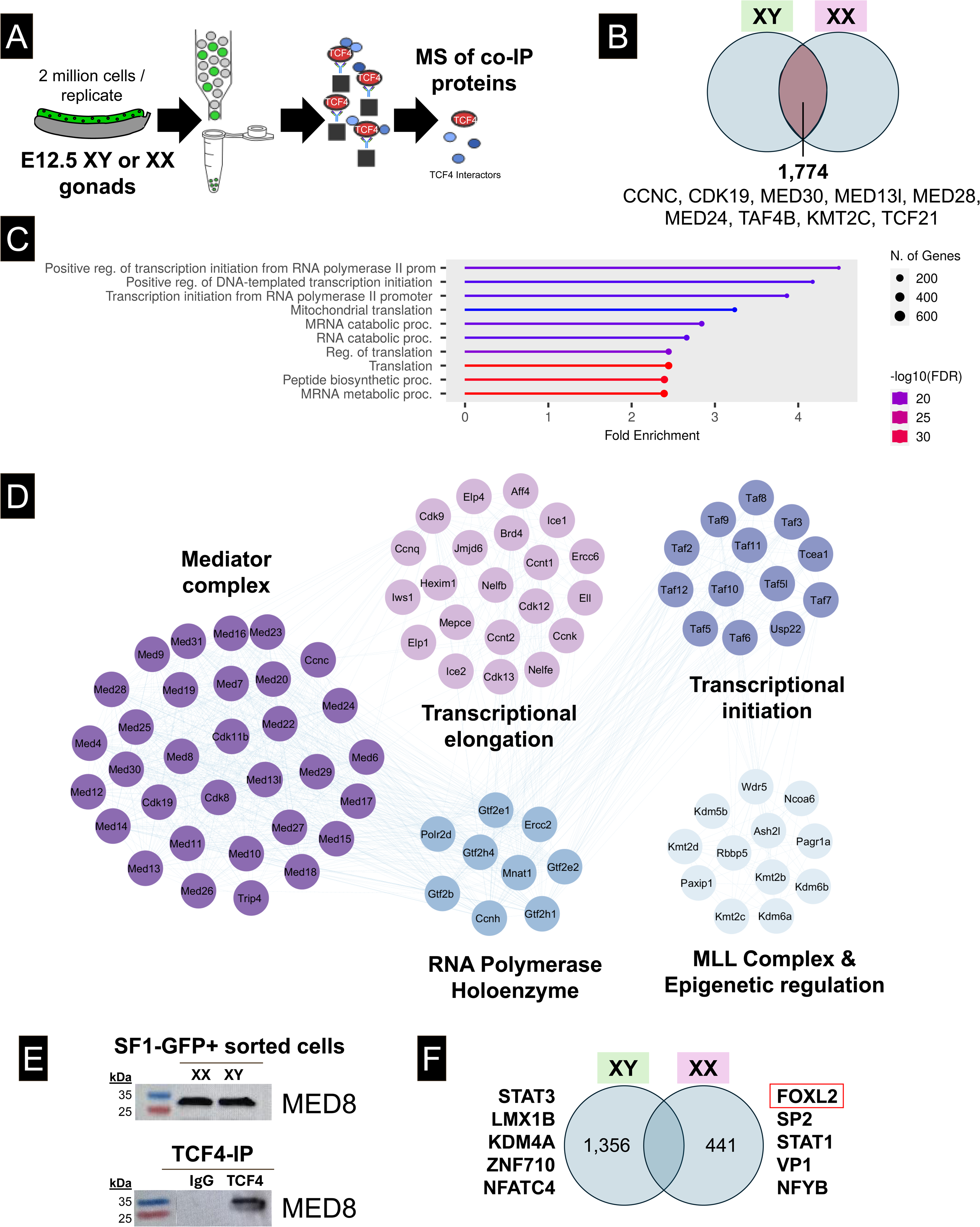
TCF4 binds core transcriptional regulators including the Mediator complex in both XX and XY sorted cells. A) *Nr5a1-Gfp* cells were sorted, immunoprecipitated for TCF4, and sent for mass spectrometry analysis to identify TCF4 binding partners. B) In total, 1,774 proteins were identified as TCF4 co-binders common to XY and XX cells. C) Proteins bound by TCF4 were enriched for processes such as positive transcription initiation and regulation of transcription, suggesting that TCF4 is a pro-transcriptional regulator in the gonad. D) Several protein complexes were identified in our mass spectrometry dataset in both XY and XX cells. Notably, the Mediator complex, as well as members of transcriptional elongation, the RNA polymerase holoenzyme, the MLL complex, and transcriptional initiation machinery. Proteins organized in a circle are known complex proteins, and lines indicate interactions. E) XY and XX *Nr5a1-Gfp* cells were sorted and western blot was performed to show the presence of Mediator 8 in somatic gonadal cells (20 µg input). Presence of Med8 was also shown in the western blot of IP’d TCF4 from XY cells (2 million cells / IP). **F**) Additionally, TCF4 co-precipitated with sex-specific proteins, including FOXL2, STAT1, and NFYB in XX cells and STAT3, LMX1B, and KDM4A in XY cells. These proteins were associated with translational machinery in XY cells, and metabolic processing in XX cells.

### TCF4 regulates gonadal genes and the transcription factor *Jun* in the ovary

Finally, ChIP-sequencing was performed using chromatin from either XX or XY *Nr5a*1-GFP cells at E12.5 to identify TCF4 targets and determine if TCF4 directly regulates genes in the granulosa network. Reads were aligned to the mm10 genome assembly, and peaks were called using MACS2 software (Supplemental Table 4). Next, stringent filtration steps were imposed on the data; peaks were filtered out if they had a Q-value < 7 or did not overlap within 1kb across all three sexed replicates (Figure 5A). While this significantly reduced the number of TCF4 targets, it enhanced the robustness of the results and minimized potential biases. In total, 96 peaks and 92 genes were identified across all three XX samples and 16 peaks and 9 genes were identified in XY samples (Figure 5B) (Supplemental Table 4).

**Figure 5.**
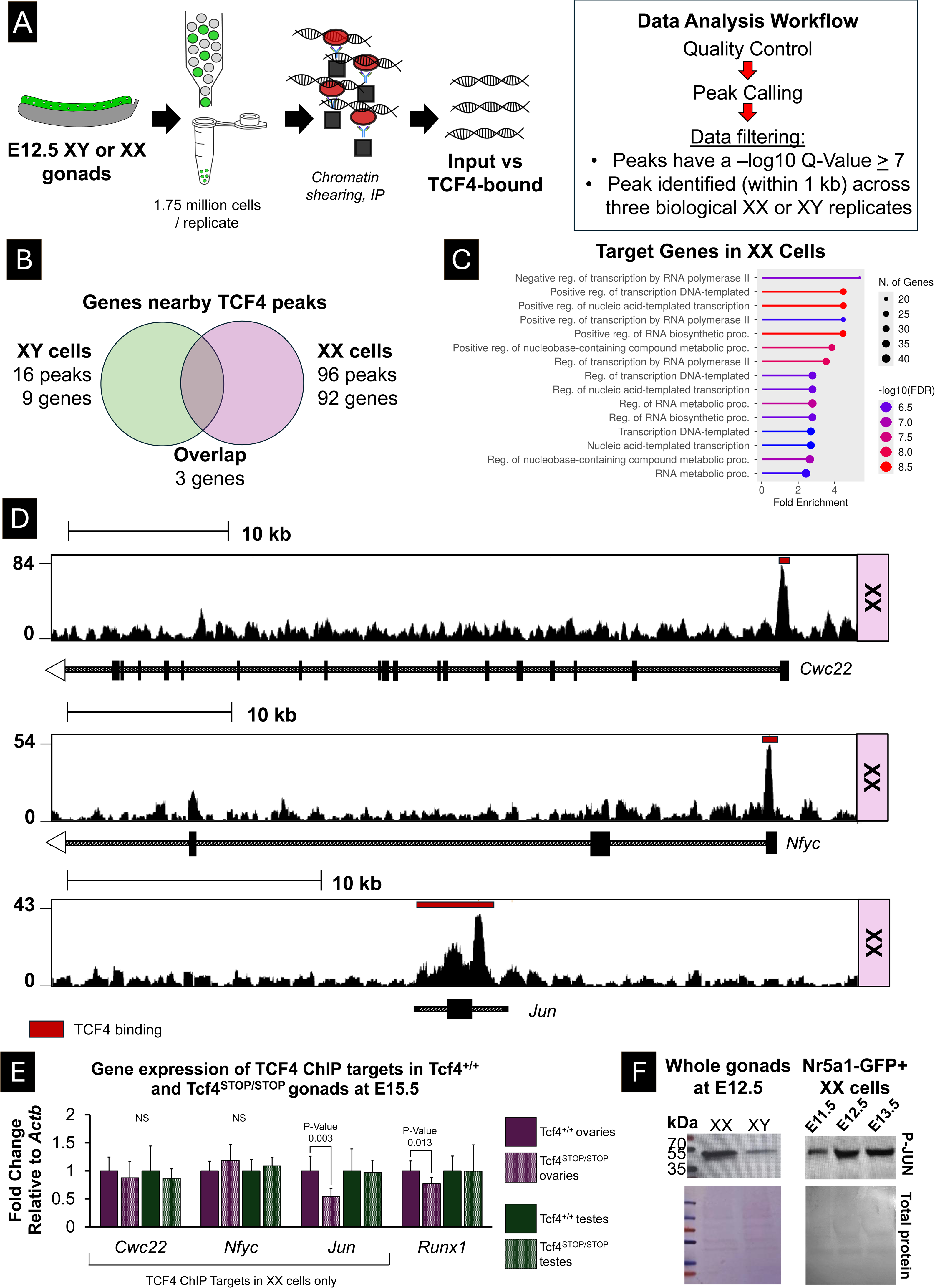
Gene targets of TCF4 include *Jun* and other transcriptional regulators. A) *Nr5a1-Gfp* cells were sorted and a ChIP-sequencing experiment was performed to identify TCF4 target genes in XX and XY chromatin. B) In total, 92 genes were identified as TCF4 targets in XX cells, while only 9 genes were identified in XY cells. C) TCF4 targets were enriched for processes including negative and positive regulation of gene transcription and RNA metabolism. D) TCF4 directly binds to the first exon of the splicing factor, *Cwc22,* the transcription factor, *Nfyc,* and across the gene body of the transcription factor *Jun.* E) *Jun,* a TCF4 ChIP target identified in XX cells, and *Runx1* expression is decreased in *Tcf4^STOP/STOP^* ovaries relative to *Tcf4^+/+^* ovaries from wildtype littermates. In contrast, *Jun* was not identified as a TCF4 ChIP target in XY cells and was not dysregulated in *Tcf4^STOP/STOP^* mutant testes. Gene expression was normalized to the housekeeping gene *Actb.* Expression is displayed as log_2_-fold change; statistics were calculated based on ΔCT value comparing wildtype and mutant gonads of the same sex. F) The active, phosphorylated form of JUN (P-JUN) is more abundant in E12.5 ovaries compared to E12.5 testes. By E15.5, P-JUN is less abundant in the embryonic ovary than testis. Additionally, P-JUN is present in sorted XX gonadal *Nr5a1-Gfp* cells across early gonad development (E10.5-E13.5) (approximately 200,000 cells per lane). Total protein stain showed similar loading of protein across the different samples.

Given that TCF4 inactivation negatively impacted ovary development, we focused on XX target genes for downstream validation. TCF4 target genes were enriched for processes in regulation of transcription and RNA metabolism (Figure 5C). TCF4 targets unique to XX cells included *Cwc22,* a splicing protein, and *Nfyc* and *Jun,* two transcription factors (Figure 5D). We measured the expression of the XX ChIP target genes *Cwc22, Nfyc*, and *Jun* in the *Tcf4^STOP/STOP^* mutant gonads relative to *Tcf4^+/+^* wildtype gonads. While the TCF4 putative targets *Cwc22* and *Nfyc* were not affected in mutants, we observed that expression of *Jun* as well as the granulosa-specific transcription factor *Runx1* were significantly reduced in *Tcf4^STOP/STOP^* mutant ovaries compared to wildtype *Tcf4^+/+^* ovaries (P < 0.05) (Figure 5E). In contrast, the expression of *Jun -* which was not identified as a TCF4 target in XY cells - and *Runx1* were not dysregulated in *Tcf4^+/+^* mutant testes relative to *Tcf4^+/+^* wildtype testes.

Although TCF4 regulates *Jun* expression in the ovary, it is unclear whether *Jun* plays a role in ovary development. Transcriptome data and single-cell RNA sequencing show that *Jun* expression becomes higher in the granulosa lineage than in the Sertoli cell lineage shortly after sex determination [5, 7]. Consistent with this, *Jun* was more highly expressed in *Tcf4^+/+^* ovaries than *Tcf4^+/+^* testes at E15.5 (Supplemental Figure 3B). However, JUN must be activated by phosphorylation (P-JUN) to work as a transcription factor – a process that occurs downstream of signaling cascades such as WNT signaling, which plays a pivotal role in ovary development. Thus, we measured P-JUN in whole embryonic ovaries and testes at E12.5 and in XX FACS-sorted *Nr5a1-GFP* gonadal cells across gonad development. We observed that activated P-JUN is more abundant in whole XX gonads compared to XY gonads at E12.5, and that P-JUN is present in Nr5a1-GFP XX somatic gonadal cells across sex determination (from E11.5 to E13.5) (Figure 5F). These results suggest that TCF4 promotes *Jun* expression, and that JUN activation is elevated in XX gonads during ovary development.

## Discussion

Commitment of the somatic gonad cells to the testicular Sertoli or ovarian granulosa lineage is the first essential step in gonad differentiation. In XY fetuses, *Sry* expression activates transcriptional changes that lead to Sertoli cell fate commitment and development of a testis [5, 7, 20, 42]. In contrast, the upstream regulators driving the granulosa gene regulatory network and *Foxl2* expression in XX cells remain unclear.

Across various studies, a group of genes essential for the formation of the bipotential gonad has emerged, including *Nr5a1, Gata4, Wt1, Lhx9,* and *Emx2* [8–10, 12–14]. To search for a common element in the regulation of these genes, we performed an investigation of TF binding sites found upstream of early regulators of gonad differentiation. Binding sites for bHLH TFs were enriched upstream of core gonadal genes, and were 1.5 times more common in genes associated with the ovary pathway than in genes associated with the testis pathway [9, 10, 12, 14].

Of the many bHLH factors expressed in gonadal cells, we chose to investigate whether TCF4, which has a known role in gene activation [43–45], plays a role in ovary development. TCF4 is highly expressed in gonadal somatic cells before and during sex determination. However, TCF4 expression is lost in SOX9-expressing cells, a pattern we would predict for a driver of ovarian fate.

To test whether TCF4 is involved in directing gonadal fate, we obtained a mutant mouse model for *Tcf4* in which the binding domain is disrupted, but the rest of the protein, including the AD domains that recruit p300 and transcriptional machinery, remain intact [16, 46]. In XY mutant gonads, the loss of TCF4 did not lead to any obvious disruptions in testis development. Instead, we observed higher expression of SOX9 in Sertoli cells. The fact that SOX9 binds upstream of *Tcf4* [47] and that loss of TCF4 led to elevation of SOX9 expression, suggests an antagonistic relationship between SOX9 and TCF4. In future studies, over-expression of TCF4 in SRY/SOX9-expressing Sertoli cells could determine whether TCF4 is inhibitory to Sertoli cell fate.

In XX wildtype gonads, TCF4 expression is maintained in granulosa cells in the ovary with highest abundance in the ovarian stroma. In XX *Tcf4* mutant gonads, we saw decreased expression of both the granulosa marker FOXL2 and the ovarian stromal marker, NR2F2 at the single-cell level. Surprisingly, the ratio of granulosa cells to stromal cells was increased in mutants, despite lower levels of FOXL2.

Further, defects in patterning and germ cell population were evident in some regions of mutant ovaries. These stronger effects are consistent with the hypothesis that TCF4 positively regulates the ovarian pathway.

To better understand how TCF4 functions in the gonad, we performed a MS analysis at E12.5 to identify TCF binding partners in XX and XY gonads. Based on mass spec analysis, binding partners identified in both XX and XY gonads included proteins associated with the Mediator Complex, the RNA polymerase complexes for initiation and elongation, epigenetic regulators, and transcription factors, such as FOXL2. These TCF4 binding partners were also reported in a paper showing that TCF4 binds with and recruits the Mediator complex to super enhancers to regulate neural stem cell differentiation [45]. If TCF4 plays a similar role in the gonad, it may bind to and open genes that establish gonadal fate and set the gonad on an ovary developmental pathway. Consistent with this, a key regulator of the ovarian pathway, FOXL2, was one of a handful of sex-specific binding partners in XX gonads. We speculate that TCF4 works in conjunction with other TF binding partners such as FOXL2 and TCF21, to specify granulosa and interstitial cell identity, respectively [37, 41, 48].

If TCF4 acts as a master regulator of gonadal genes, it is unclear why the phenotype in both XX and XY gonads is not more severe. This could be because of the complexity of the Mediator Complex in which loss of a single factor has a small impact, or, alternatively, because the *Tcf4* mutation leaves some functions of the TCF4 protein intact. We favor the latter interpretation, which needs further study.

If an effort to identify other upstream regulators of granulosa cell identity previously not recognized, we performed a TCF4 ChIP-sequencing analysis on E12.5 XX and XY gonads. Using stringent methods to confirm binding sites in XX gonads, we identified *Jun*, a transcription factor that is dysregulated in *Tcf4^STOP/STOP^* mutant ovaries and known to be activated downstream of WNT signaling through the planar cell polarity pathway [49–51]. Our data show that TCF4 promotes *Jun* expression, which is preferentially activated in the embryonic ovary relative to the testis at E12.5. These findings suggest that TCF4 may enhance the availability of JUN as a key mediator of WNT-driven signaling during ovary development.

In conclusion, we hypothesize that *Tcf4* is silenced in Sertoli cells to permit their differentiation in response to *Sry* expression. Our data suggest that TCF4 recruits the Mediator complex to specific chromatin regions to maintain the progenitor state of other somatic gonadal cells and partners with proteins such as FOXL2 and TCF21 to regulate expression of granulosa and stromal factors and promote ovary differentiation (Figure 6). Many other bHLH proteins are expressed in the gonad, and further investigation of this interesting class of TFs is warranted.

**Figure 6.**
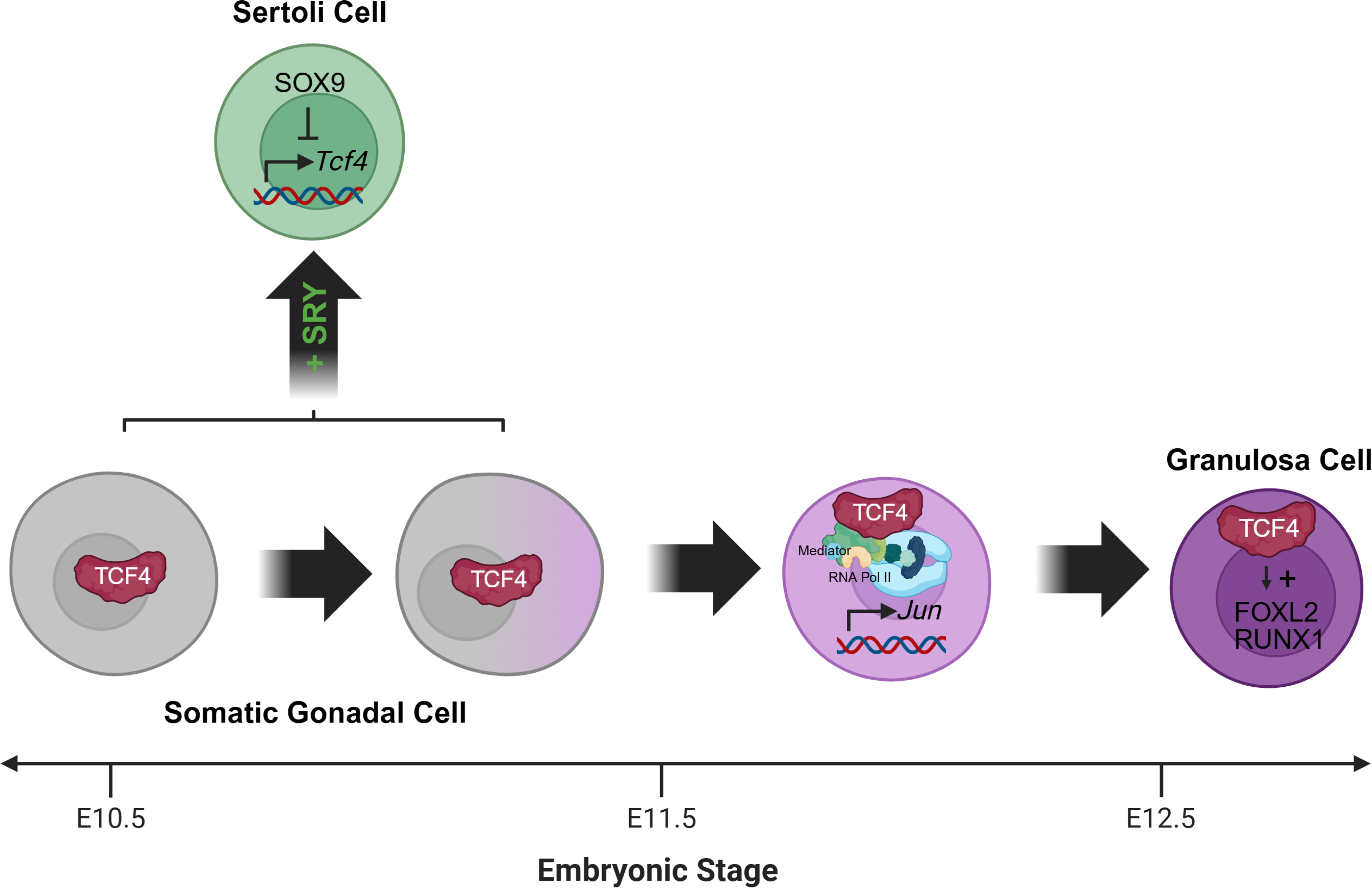
Working model for the role of TCF4 in gonad differentiation. TCF4 drives a gonadal program, likely through recruitment of the Mediator complex and epigenetic factors, that advances ovarian fate and is specifically silenced in Sertoli supporting cells as these pathways diverge. Figure created with Biorender.

## Materials & Methods

### Mouse strains and lines

All mice were housed in accordance with the National Institutes of Health guidelines, and experimental procedures were approved by the Duke University Medical Center Institutional Animal Care and Use Committee. For wildtype matings, CD-1 mice were obtained from Jackson Laboratories. The Tg(*Nr5a1-GFP)* (*Sf1-eGFP)* mice have been previously described [40]. The Tg(*Runx1-EGFP)*E1184Gsat/mmucd (*Runx1-eGFP)* mice were a gift from Humphrey Yao (MMRRC 010771-UCD) [52]. The *Tcf4^STOP/+^* mutant mouse line was a gift from Ben Philpot at UNC where they have been previously characterized [39, 53], and was maintained on a C57BL/6J (B6) background in our colony.

### Timed matings

To obtain embryos at specific embryonic stages, females were placed in timed mating cages with studs. Female mice were checked daily for the presence of a vaginal plug, which was considered embryonic day 0.5. Female CD-1 mice were used for all experiments except those involving *Tcf4^STOP/+^ mice*. For *Tcf4^STOP/+^* matings, *Tcf4^STOP/+^* B6 studs were crossed with *Tcf4^STOP/+^* B6 females.

*Organ collection and genotyping.* Gonad-mesonephric complexes were microdissected into PBS and used for downstream applications. Tail somites were counted to ensure mice were at the appropriate developmental stage. Tails or limbs were recovered from each fetus and digested in 150 mM NaOH at 95°C for 15 minutes and neutralized with 1M Tris HCl (pH 7.4) for genotyping. Fetuses were sexed by genotyping for the *Utx* and *Uty* genes using the following three primers (5’➔3’): UTX F1: TCATGTCCATCAGGTGATGG, UTY F2: CAATGTGGACCATGACATTG, UTX/Y R: ATGGACACAGACATTGATGG. Mice from the *Tcf4^STOP/+^* breeding scheme were genotyped for the wildtype allele (primer sequences 5’ ➔ 3’ F: GCACTTCAGGGATCGCTTA, R: CCGCCCTAATTGTTCAAAGAG) or transgenic knock-in allele (primer sequences 5’ ➔ 3’: F: GCTGATCCGGAACCCTTAAGC, with the TCF4 WT R primer). PCR conditions have been previously reported [53, 54]. In the case of *Nr5a1-GFP* mice, fluorescent signal was strong enough to observe with the NIGHTSEA fluorescence detection system on the dissection microscope (NIGHTSEA, Lexington, MA), bypassing the need for genotyping.

### Whole mount Immunofluorescence

Embryonic gonads were fixed for 1 hour at room temperature (RT) or overnight (O/N) at 4°C in 4% paraformaldehyde while rocking. Samples were rinsed 3 times in PBS and then passed through an increasing methanol gradient in PBS (25%, 50%, 75%, 100%). Gonads were stored in 100% methanol O/N or longer, as needed. Samples were rehydrated stepwise into PBS and rinsed 3 times in PBS. Late-stage gonads (E15.5) were incubated in 2% Triton X-100 in PBS for 1 hour at RT to increase permeabilization. Early and late-stage gonads were then incubated in blocking buffer (0.1% Triton X-100 for E10.5-E13.5 or 1% Triton X-100 for E15.5 and later, with 3% BSA, and 10% fetal bovine serum or horse serum [HS] in PBS) for 1 hour. Tissue was incubated with primary antibody in blocking solution O/N. Antibody details and concentrations are listed in Table 1. The following day gonads were rinsed in either 0.1% (E10.5-E14.5) or 1% Triton X-100 (E15.5 and later) in PBS and incubated with secondary antibodies (1:500) in blocking buffer O/N. Gonads were rinsed 3 times for 10 minutes in PBS and then mounted in DABCO mounting media. Images were captured with a Leica TCS SP8 confocal microscope using the associated Leica software (Leica, Wetzlar Germany). Images were processed using FIJI (ImageJ).

**Table 1.** Key Resources.

| Reagent type (species) or resource | Designation | Source or reference | Identifiers | Additional information |
| --- | --- | --- | --- | --- |
| Gene (*M. musculus*) | Transcription factor 4 (Tcf4) | GenBank | MGI:98506 |  |
| Strain, strain background (*Mus musculus*) | CrI:CD1(ICR) | Charles River | Strain code: 022 |  |
| Strain, strain background (*M. musculus*) | C57BL/6J | Jackson Laboratory | Stock #:000664 |  |
| Genetic reagent (*M. musculus*) | Tg(Nr5a1/EGFP)1Klp | PMID: 12351700 | MGI:5493455 |  |
| Genetic reagent (*M. musculus*) | Tg(Tcf4<sup>STOP/+</sup>) | PMID: 32765228 |  |  |
| Antibody | Smooth muscle alpha actin (aSMA) (FITC-conjugated mouse monoclonal) | Sigma-Aldrich | F3777 | 1:500 |
| Antibody | Endostatin/Collagen 18 (goat polyclonal) | R&D Systems | AF570 | 1:500 |
| Antibody | FOXL2 (rabbit polyclonal) | Gift from Dagmar Wilhelm |  | 1:250 |
| Antibody | GATA4 (mouse monoclonal IgG<sub>2α</sub>) | Santa Cruz Biotechnology | Sc-25310 | 1:250 |
| Antibody | GFP (chicken polyclonal) | Abcam | Ab13970 | 1:1000 |
| Antibody | Phospho-c-Jun (Ser73) (rabbit monoclonal) | CellSignaling Technology | (D47G9) | 1:1000 |
| Antibody | Phospho-JUN (Ser73) | Proteintech | 4A18 | 1:1000 |
| Antibody | MED8 (rabbit polyclonal) | Proteintech | 12182-1-AP | 1:1000 |
| Antibody | NR2F2 (mouse monoclonal IgG<sub>2α</sub>) | Abcam | Ab41859 | 1:200 |
| Antibody | NR5A2 (goat polyclonal) | Santa Cruz Biotechnology | Sc-21132 | 1:500 |
| Antibody | SOX9 (goat polyclonal) | R&D Systems | AF3075 | 1:500 |
| Antibody | TCF4 (rabbit monoclonal) | Abcam | Ab217668 | 1:500 |
| Antibody | TRA98/GCNA1 (rat monoclonal) | Abcam | Ab82527 | 1:500 |
| Antibody | AF488 Anti-Mouse (donkey polyclonal) | Jackson ImmunoResearch | 715-545-150 | 1:500 |
| Antibody | AF488 anti-Rat (donkey polyclonal) | Life Technologies | A-21208 | 1:500 |
| Antibody | AF488 anti-Chicken (donkey polyclonal) | Jackson ImmunoResearch | 703-545-155 | 1:500 |
| Antibody | Cy3 anti-Goat (donkey polyclonal) | Jackson ImmunoResearch | 705-165-147 | 1:500 |
| Antibody | Cy3 anti-Mouse (donkey polyclonal) | Jackson ImmunoResearch | 715-165-150 | 1:500 |
| Antibody | AF647 anti-Rabbit (donkey polyclonal) | Jackson ImmunoResearch | 711-605-152 | 1:500 |

### Mapping HLH, SOX, FOX, and COUP binding sites

HLH motif file was generated for the E-box motif (CANNTG) using the “seq2profile” command from HOMER. HLH sites genome wide were identified using the “scanMotifGenomeWide” tool against the mm9 genome [55]. The resulting bed file from this motif scan, as well as bed files from SOX9 and FOXL2 ChIP-seq studies [47, 56], were processed using the tool “annotatePeaks” to identify nearest gene transcription start sites to each genomic location present in the bed files. Next, gene lists unique to XX and XY supporting and interstitial cells were filtered by their fold-change between E11.5 and E13.5. Genes that showed a 1.15-fold change and that were statistically significant (P < 0.05) were defined as ‘Increasing’ or ‘Decreasing’ in the supporting and interstitial genes and output as a csv file (Supplemental Table 1). Annotated bed files were converted to csv and uploaded to R studio alongside the gene lists. Gene lists and motif/ChIP sites were compared for overlap using the function “inner_join” based on Gene Symbol. HLH binding sites were searched for within 1 kilobase of the gene of interest.

### Fluorescence Activated Cell Sorting (FACS) Analysis

For cell analysis of E15.5 *Tcf4^+/+^* or *Tcf4^STOP/STOP^* gonads, gonad pairs from single fetuses were digested in TrypLE (Gibco, Thermo Fisher, Waltham MA) with 0.1% collagenase for 10 minutes at 37°C. TrypLE was removed from samples and samples were then resuspended in 3% BSA in PBS and filtered through a 50-um filter. Samples were spun down at 600 g for 10 minutes at 4°C and then fixed in 4% PFA with 0.1% Triton X-100. After a 10-minute incubation, samples were spun down again at 600 g for 10 minutes at 4°C. Samples were left in 3% BSA in PBS at 4°C for up to ten days before primary antibody staining. Cells were recovered, spun down at 600g for 10 minutes at 4°C, and then incubated in blocking buffer (10% HS with 0.1% Triton X-100 in PBS) O/N with an antibody to mark interstitial cells (NR2F2) or supporting cells (SOX9/FOXL2 antibody depending on the sex of the sample) (Antibody details in Table 1). The following day samples were spun down at 600 g for 10 minutes at 4°C and placed in blocking buffer with secondary antibody (Donkey anti-Mouse Cy3, donkey anti-rabbit AF647, or donkey anti-goat AF647) for 1 hour at RT. Samples were spun down and rinsed in 3% BSA in PBS twice before analysis. For FACS analysis, wildtype and mutant gonad suspensions were run on an BD LSRFortessa™ Analyzer in a single session. The area of the forward side scatter (FSC-A) and side scatter area (SSC-A) were plotted to isolate cells and remove debris, and the FSC-A and FSC-H were plotted to obtain single cells. Gating conditions were established using unstained gonad cells, and single fluorophore controls (AF657 and Cy3). Voltage settings were as follows: SSC 246, FSC 360, APC 571, PE 505. Next, 5,000 cells from gonad pairs of individual, genotyped *Tcf4^WT/WT^ and Tcf4^STOP/STOP^* fetuses were recorded. Gating was set at a signal of > 10^2.5^ for the 647 fluorophore and > 10^3.1^ for the Cy3 fluorophore. Cells with fluorescence signal greater than these gating values were identified as ‘supporting’ or ‘interstitial’ cells. The ratio of supporting: interstitial cells was then calculated and compared between wildtype and mutant conditions using a two-tailed t-test. The geometric mean, which gives the average fluorescence on a single-cell level in heterogenous populations, was also determined and compared between wildtype and mutant conditions using a two-tailed t-test.

### Cell sorting

For cell sorting of *Nr5a1-GFP* gonads at E12.5, fetuses were sexed by PCR, and the gonads were dissected, pooled by sex, and dissociated for 10 minutes in TrypLE at 37°C. After digestion, TryPLE was carefully removed, and gonads were resuspended in 10% HS in PBS. The cell suspensions were filtered through a 50-micron filter and sorted on either a BD FACSAria or Astrios Cell Sorter. GFP expression was used to sort somatic cell types, with gating set using a GFP-negative control sample by a Duke Flow Cytometry Core specialist. Cells were recovered and prepared in one of two ways for downstream analysis. For downstream IP-MS and western blot analysis, samples were spun down at 3500 g for 10 minutes at 4°C, snap-frozen, and stored at −80°C. For ChIP-sequencing, cells were first centrifuged at 500 g for 10 minutes at 4°C, rinsed twice in PBS +/+, and fixed for 10 minutes at RT in freshly prepared 1% formaldehyde. Cross-linking was quenched with 2.5 M glycine and incubated for 5 minutes at 4°C. Cells were then rinsed twice in ice cold PBS and lysed in cell lysis buffer (10 mM Tris HCl pH 7.5, 10 mM NaCl, 3 mM MgCl_2_, 0.5% NP-40, 1 mM DTT, 1 mM PMSF, 1x Protease inhibitor) for 10 minutes at 4°C. Nuclear pellets were spun down at 500 g for 5 minutes at 4°C, the supernatant was removed, and samples were stored at −80°C for later pooling *IP-MS.* Frozen sorted somatic gonadal cells were pooled into three replicates of 4 million cells (± 10%). Cell pellets were lysed in a Pierce™ IP Lysis Buffer (Thermo Fisher) containing 1 mM PMSF and 1x Halt™ Protease Inhibitor Cocktail (Thermo Fisher), incubated on ice for 30 minutes, and centrifuged at 14,000 g for 10 minutes at 4°C. Meanwhile, 50 µl of Protein G Dynabeads (Invitrogen, Waltham MA) were rinsed 3 times in PBS with 0.05% Tween-20 using a magnet bar. Beads were suspended in 237.5 µl of Conjugation buffer (20 m HEPES in PBS), and 5 µg of IgG or TCF4 antibody (Abcam, NCI-R159-6) was crosslinked to the beads using 12.5 µl of 100 mM BS3. Beads were gently vortexed and incubated for 30 minutes in crosslinking buffer. After incubation, beads were resuspended in 250 µl of Conjugation buffer and quenched with 12.5 µl of 1 M Tris HCl (pH 7.5). Beads were incubated for 15 minutes at RT and rinsed 3 times in PBS with 0.05% Tween-20. The extracted protein was then split between IgG or TCF4-conjugated beads and incubated O/N on an end-over-end rotator at 4°C. The following day beads were rinsed in dilution buffer (10 mM Tris HCl pH 7.5 with 150 mM NaCl and 0.5 mM EDTA), dilution buffer with 1% Triton X-100, dilution buffer with 1% Triton X-100 and 360 mM NaCl, and finally in dilution buffer. Proteins were eluted at 80°C for 10 minutes in 25 mM Tris, 50 mM NaCl, and 1% SDS and submitted to the Duke Proteomics Core for MS analysis.

### Quantitative LC-MS

Samples were spiked with 1 or 2 pmol bovine casein as an internal quality control standard. The samples were brought to a final concentration of 5% SDS and reduced for 15 minutes at 80°C. After reduction, the samples were alkylated with 20 mM iodoacetamide for 30 minutes at RT, then supplemented with 1.2% phosphoric acid and 416 µl of S-Trap (Protifi, Firport NY) binding buffer (90% MeOH/100 mM TEAB). Proteins were trapped on the S-Trap microcartridge, digested with 20 ng/µl of trypsin (Promega, Madison WI) for 1 hour at 47 °C, and eluted sequentially with 50 mM TEAB, followed by 0.2% formic acid (FA) and 50% aceconitrile/0.2% FA. All samples were lyophilized and resuspended in 1% TFA/2% acetonitrile with 12.5 fmol/µl of yeast alcohol dehydrogenase. A study pool sample was created by combining equal volumes of each sample. Quantitative LC/MS was performed on an EvoSep One UPLC coupled to a Thermo Orbitrap Astral high-resolution, accurate-mass tandem mass spectrometer (Thermo Fisher). Each sample was eluted from the EvoTip onto a 1.5 µm EvoSep 150 µm ID x 1 5cm performance EvoSep column using the SPD30 gradient at 55°C. Data collection on the Orbitrap Astral mass spectrophotometer was performed in a data-independent acquisition (DIA) mode of acquisition with a resolution of 240,000 for full MS scans in the m/z range of 380-980. An HCD collision energy setting of 27% was used for all MS2 scans.

### Quantitative Proteomics Data Analysis

Raw data were imported into Spectronaut (Biognosis, Zurich Switzerland) and individual LC-MS data files were aligned based on the accurate mass and retention time of detected precursor and fragment ions. Relative peptide abundance was measured based on MS2 fragment ions of selected ion chromatograms of the aligned features across all runs. The MS/MS data was searched against a SwissProt M. musculus database and an equal number of reversed-sequence “decoys” for false discovery rate determination. A library free Direct DIA+ approach within Spectronaut was used to perform the database searches. Database search parameters included a fixed modification on Cys (carbamidomethyl), with variable modification on Met (oxidation) and Lys (biotin). Full trypsin enzyme rules were applied, along with 10ppm mass tolerances on precursor ions and 20ppm on product ion. Spectral annotation was set at a maximum 1% peptide false discovery rate, based on q-value calculations. Peptide homology was addressed using razor rules, where a peptide that matches to multiple different proteins is exclusively assigned to the protein with the most identified peptides. Protein homology was handled by grouping proteins that had the same set of peptides to account for their identification. Fold-changes were calculated between the IgG and TCF4 group using a two-tailed heteroscedastic t-test on log2-transformed data. The P-Value was rounded to two decimals points. Proteins were enriched TCF4 binding partners if they met the criteria of being at least 1.5-fold higher than the IgG value with a P-value ≤ 0.05.

### Western blot analysis

Protein was extracted from *Nr5a1-GFP* cells using Pierce™ RIPA Lysis Buffer (Thermo Fisher) with 1 mM of PMSF and 1x Halt Protease and Phosphatase inhibitor cocktail (Thermo Fisher). Protein isolates from *Nr5a1-Gfp* cells were quantified using a BCA assay. For the western blot, equivalents amount of protein [200,000 sorted cells worth of protein extract for the P-JUN western blot in *Nr5a1-GFP* cells, 20 µg of measured protein extract for the western blot measuring P-JUN and MED8 in whole gonads, or 30 µl of eluant from the IgG or TCF4 IP (approximately half the total eluant from 4 million cell IP reaction split between IgG or TCF4 antibody)] was reduced with 5x Laemmli buffer and 1 mM DTT at 95°C for 5 minutes. Samples were loaded onto a mini-PROTEAN TGX precast gel (Bio-Rad, Hercules, CA) and proteins were separated via SDS-PAGE at a constant voltage of 100 V for 90 minutes in running buffer (25 mM Tris, 192 mM glycine, 0.1% SDS). Protein was transferred using a Trans-Blot Turbo Transfer System (Bio-Rad), following the Manufacturer’s Instructions. Membrane was blocked for 1 hour at 5% milk in 0.1% Tris-Buffered saline with 0.1% Tween-20 (TBST) and incubated O/N with a polyclonal rabbit MED8, monoclonal rabbit P-JUN, or monoclonal rabbit TCF4 antibody. Blot was rinsed three times in 0.1% TBST and then put in blocking buffer with goat anti-rabbit horseradish peroxidase (HRP) conjugated secondary antibody at a 1:10,000 dilution for 1 hour at RT. Blot was rinsed three times in 0.1% TBST and developed in Super Signal West Pico Chemiluminescent Substrate (Thermo Fisher) for 5 minutes before visualization on an Amersham Imager 600. For non-IP western blots, blots were stripped in Restore PLUS Western Blot Stripping buffer for 15 minutes at RT. Blots were blocked for 1 hour in 5% milk in 0.1% TBST again and then either stained with Colloidal Gold Total Protein Stain (Bio-Rad) or incubated with a polyclonal goat anti-GAPDH antibody (1:5,000) O/N. The following day, the blot was rinsed three times in 0.1% TBST and incubated in a donkey anti-goat horseradish peroxidase (HRP) conjugated secondary antibody. Development and imaging were repeated as previously described.

### ChIP Sequencing

Nuclear pellets from approximately 1.75 million (± 10%) sorted *Nr5a*1-GFP cells were pooled and placed in a MNase reaction buffer (10mM Tris HCl pH 7.5, 10mM NaCl, 3 mM MgCl2, 1 mM Cacl2, 0.2% NP-40, 1 mM DTT, 1mM PMSF, 1x protease inhibitor). Nuclei were centrifuged at 500 g for 5 minutes at 4°C and placed in 200 µl of fresh MNAse reaction buffer. Next, 55 units of MNase was added to each tube and nuclear pellets were incubated for 15 minutes at 37°C with vortexing every five minutes. Enzymatic DNA fragmentation was stopped by adding 20 µl of 0.5 M EDTA. Samples were centrifuged at 9400 g for 10 minutes at 4°C and placed in 75 µl of ChIP sonication buffer (50 mM Tris HCl pH 7.9, 10 mM EDTA, 1% SDS, 1mM PMSF, 1x protease inhibitor). Samples were left on ice for 10 minutes and 75 µl of ChIP dilution buffer (16.7 mM Tris HCl pH 7.9, 1.1% Triton X-100, 167 mM NaCl, 1.2 mM EDTA, 1 mM PMSF). DNA was sonicated four times for 20 seconds with 30 seconds of rest. Samples were centrifuged at 13000g for 10 minutes at 4°C. Next, 600 µl of ChIP dilution buffer was added to samples, 2% of the sample volume was collected for input, and 5 µg of TCF4 antibody was added for overnight incubation at 4°C on an end-over-end rotator. The following day, 15 µl of Dynabeads Protein G magnetic beads were added to each tube. Samples were incubated for two hours on an end-over-end rotator. Using a magnetic bar, the supernatant was removed and beads were sequentially rinsed for 10 minutes in low salt buffer (20 mM Tris HCl pH 7.9, 100 mM NaCl, 2 mM EDTA, 0.1% SDS, 1% Triton X-100), high salt buffer (20 mM Tris HCl pH 7.9, 500 mM NaCl, 2 mM EDTA, 0.1% SDS, 1% Triton X-100), LiCl Buffer (10 mM Tris HCl pH 7.9, 250 mM LiCl, 1% NP-40, 1% deoxycholate acid, 1 mM EDTA) and TE buffer. Next, 200 µl of elution buffer (1% SDS, 0.1 M NaHCO_3_) was added and samples were left at 65°C for 40 minutes to elute DNA. At this point, input DNA was thawed, brought to 200 µl with elution buffer, and processed in parallel with samples. Samples were recovered, 8 µl of 5 M NaCl (per 200 µl of elution buffer) was added to samples, and samples were left at 55°C overnight to de-crosslink the DNA. The next day, 14 µl of Proteinase K solution was added to each sample (4 µl of 0.5 M EDTA, 8 µl of 1 M Tris HCl pH 6.8, 2 µl of 20 mg/ml Proteinase K) before a 2-hour incubation at 55°C, and DNA was eluted using the Qiagen MinElute on column PCR Purification kit (Qiagen, Hilden, Germany). DNA was eluted in 10 ul of TE buffer.

### ChIP sequencing data analysis

Samples were submitted to BGI Genomics for sequencing and analysis. Briefly, immunoprecipitated DNA was purified and checked for quality using a Fragment Analyzer. Sequencing libraries were prepared by adding ligating sequencing adaptors and sequenced on the DNBSEQ platform to generate 50 bp paired-end reads. After adapter trimming and removal of low-quality reads, an average of approximately 25 million high-quality reads were obtained per sample. Clean reads were aligned to the reference genome mm10 using SOAPaligner/SOAP2 software with two mismatches. Only uniquely mapped reads with a mapping quality score (MAPQ) greater than 20 were retained. Peak calling was performed using MACS1.4.2 with a Q-value cutoff of < 0.05 relative to the input control. Individual peak lists were generated for the six replicates (three sexed replicates). We then selected high-confidence peaks (Q-value < 10^-7) and designed a replicate-matching algorithm to only select peaks that were consistently observed across the three sexed replicates. Specifically, we computed clusters of peaks such that each cluster a) contains at least one peak from each replicate, and b) each peak in the cluster is within 1 kb of at least one peak from another replicate. This was achieved by constructing a tri-partite graph across three replicates where nodes are peaks, edges connect cross-replicate peaks within 1 kb and then identifying connected graph sub-components that span all replicates. Peaks not assigned to any cluster were discarded. The Python implementation of the matching algorithm will be made available to the research community as an open-source package on Github. The peaks called after stringent filtration as well as the peak lists for each individual replicate are listed in Supplementary Table 4. Individual bigWig files are available for download at DOI 10.17605/OSF.IO/H6D2M. Raw FASTQ files will be deposited in GEO and made publicly available upon publication.

### qPCR analysis

Tissue was dissected and run through a QiaShredder column in Buffer RLT (Qiagen). RNA was isolated using the MinElute RNA Kit (Qiagen), including an on-column DNAse digestion. The RNA was quantified using a Nanodrop and the purity was assessed using A260/280 values. Next, 50 ng of RNA was reverse-transcribed into cDNA using the Verso cDNA Synthesis kit. Parallel reactions were run without enzyme to assess genomic contamination. Reverse transcription was performed following manufacturers’ instructions. Real-time PCR was then performed using a QuantStudio 6 Flex Real-Time System (ThermoFisher) with SsoAdvanced Universal SyberGreen Supermix (BioRad). PCR conditions were: activation, 95°C for 15 min; 40 cycles of 95°C for 15 s; 60°C for 60 s. Primers were synthesized by Integrated DNA Technologies (Primer sequences in Table 2). Technical triplicates were used to generate the cycle threshold for each gene in each sample. Melt curve analysis was performed to verify PCR product specificity. ΔCT values were calculated as the difference between the cycle threshold (CT) of the gene of interest and the housekeeping gene (*Actb)*. The housekeeping gene *Actb* is highly and stably expressed during early gonad development [5, 7]. Statistics were calculated using the ΔCT values. For visualization, fold change was calculated relative to the reference gene. Standard error was calculated by propagating the standard deviations of the gene of interest and housekeeping gene using the formula SE_ΔΔCT_= √[(SD_GOI/ √n)^2^)+(SD_HK/ √n)^2^]. Fold change was plotted as 2^-ΔΔCT^ ± propagated error.

**Table 2.**
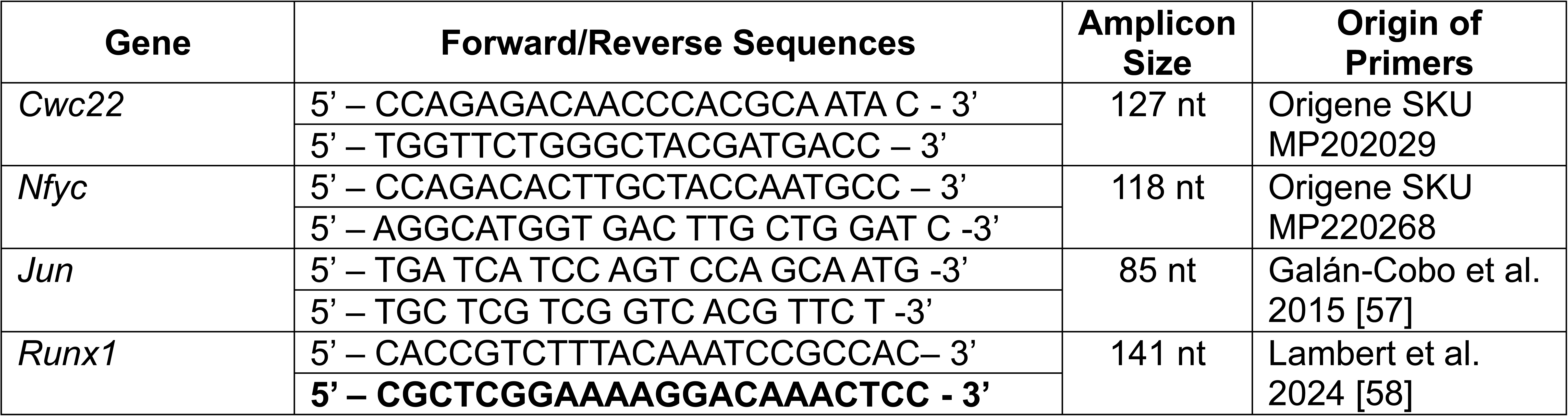
qPCR Primer Sequences.

## Acknowledgments

The authors would like to thank Ben Philpot at UNC Chapel Hill for generously providing the Tcf4^STOP/+^mouse line. The authors thank the past and present members of the Capel Lab for their support, especially Megan Harward for providing animal care and maintenance and Talia Hatkevich for assistance in FACS analysis. Finally, the authors would like to thank Erik Soderblom, Tricia Ho, and Greg Waitt for their technical assistance with mass spec sample analysis and interpretation. The authors also appreciate the Staff at the Duke Light Microscopy Core Facility and the Duke Cancer Institute Flow Cytometry Shared Resource for their excellent technical support.

## Author contributions

EVO, SMD, and BC conceptualized and designed research; EVO and BC performed research; EVO, SMD, RH, and BC analyzed data; and EVO and BC wrote the manuscript. All authors edited and approved the final manuscript.

---

**Figure S1. Expression of TCF4 is maintained in gonadal granulosa and interstitial cells through late gestation.** A) At E12.5, TCF4 co-localizes with VCAM- and NR2F2-expressing interstitial cells in XY gonads. B) At E18.5, TCF4 remains restricted to the testicular interstitial cells and absent in SOX9-expressing Sertoli cells. C) At E18.5, TCF4 is broadly expressed across both granulosa and interstitial cell lineages in the oary. Scale bar represents 100 µm. Microscopy images are stitched together from individual panels.

**Figure S2. The *Tcf^STOP^* allele is unable to bind DNA.** A) The *Tcf^STOP^* allele introduces a premature STOP codon upstream of the bHLH domain in exon 18. B) Western blot analysis shows that *Tcf^STOP/+^* mice express a truncated form of TCF4 without the bHLH domain that is absent in *Tcf^+/+^* wildtype littermates.

**Figure S3. Minor defects** in *Tcf^STOP/STOP^* mutant ovaries. A) One *Tcf^STOP/STOP^* mutant ovary displayed defects in patterning, as germ cells were present in the posterior portion of the ovary (white arrowheads) but were absent in the anterior portion. B) The transcription factor *Jun* is expressed in *Tcf^+/+^* wildtype testes at about half the level of *Tcf^+/+^* wildtype ovaries.

