## Supplementary figures and images for "The basic helix-loop-helix transcription factor TCF4 recruits the Mediator Complex to activate gonadal genes and drive ovarian development"

### Supplemental Figures

**Supplemental Figure 1**

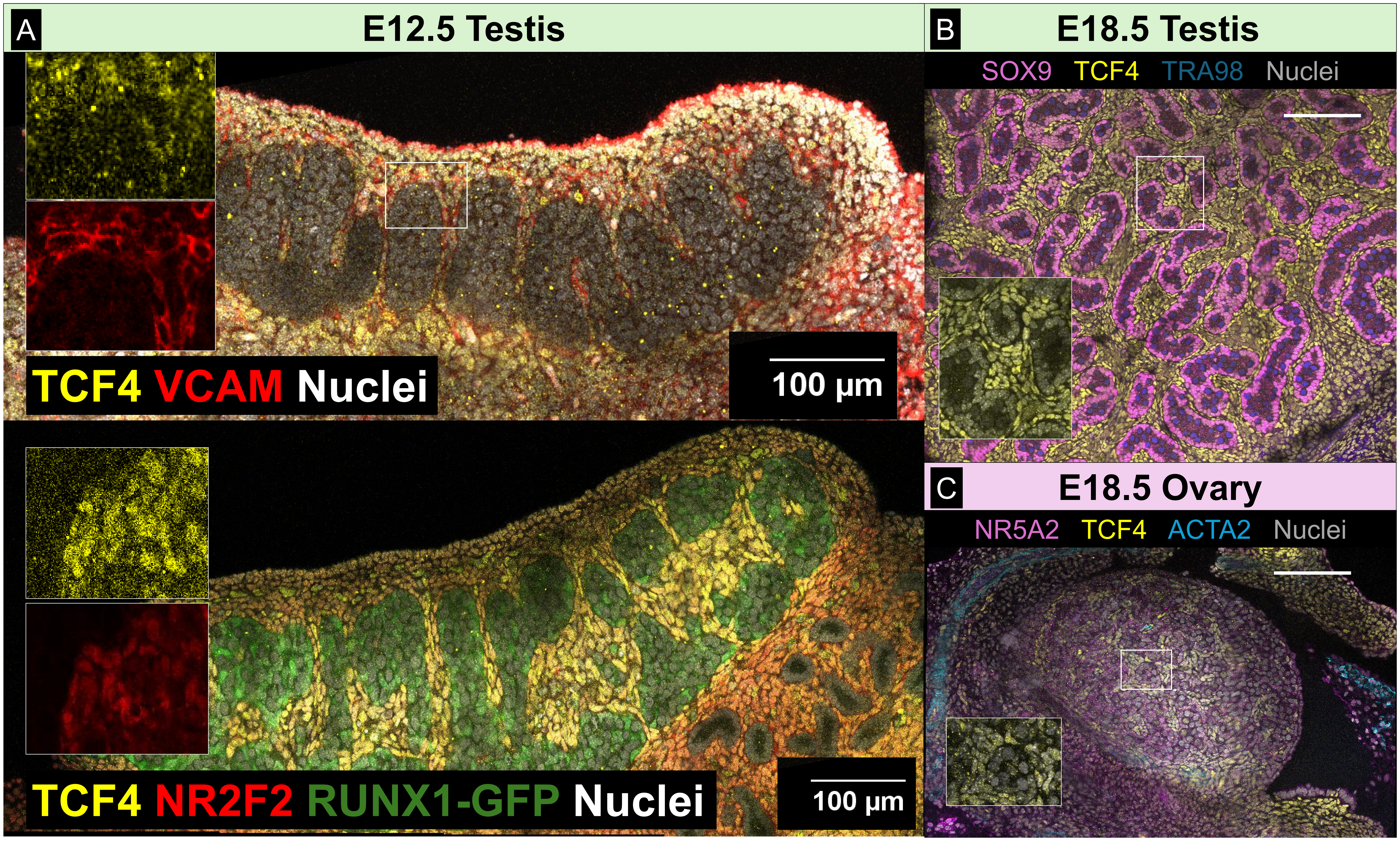

**Supplemental Figure 2**

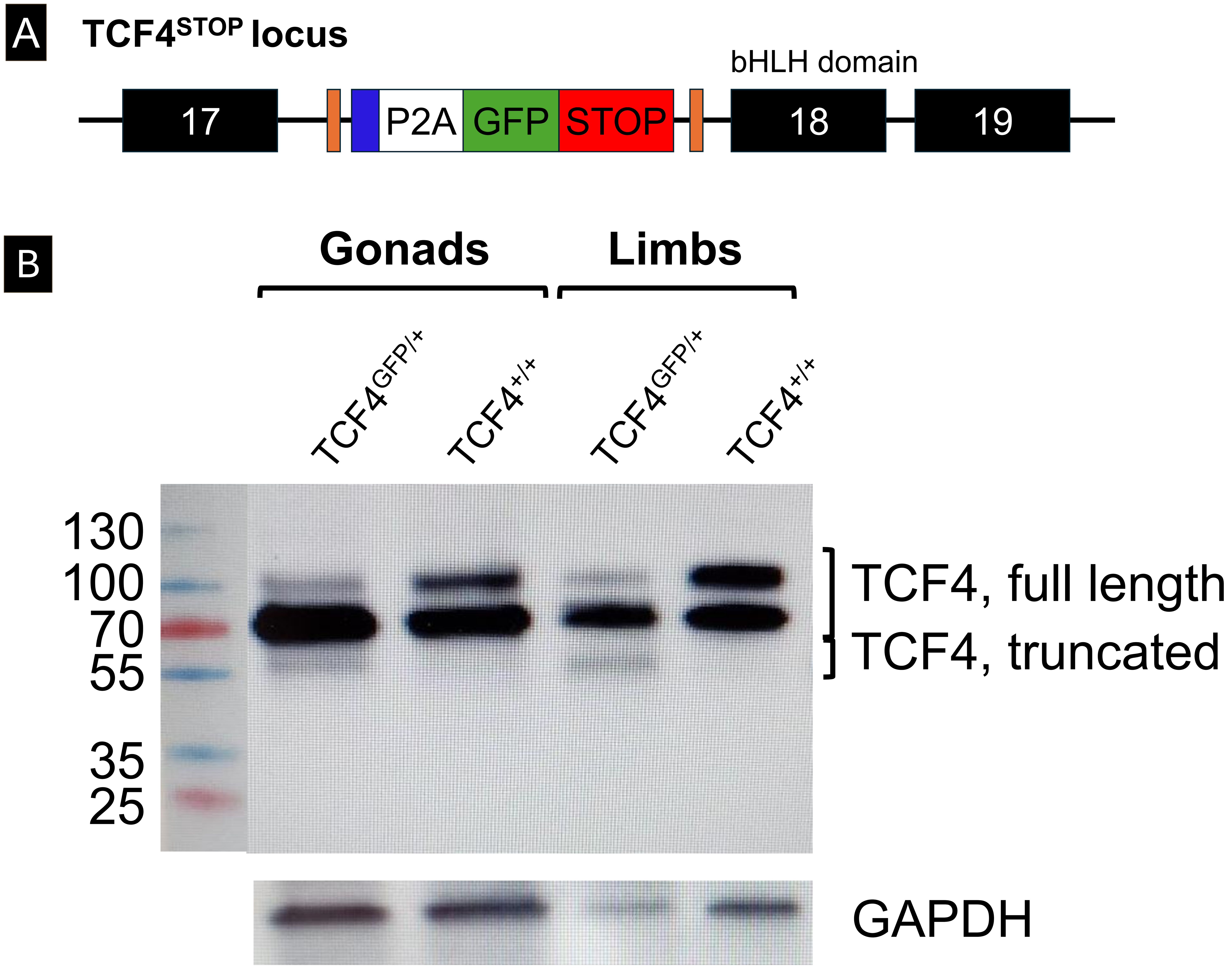

**Supplemental Figure 3**

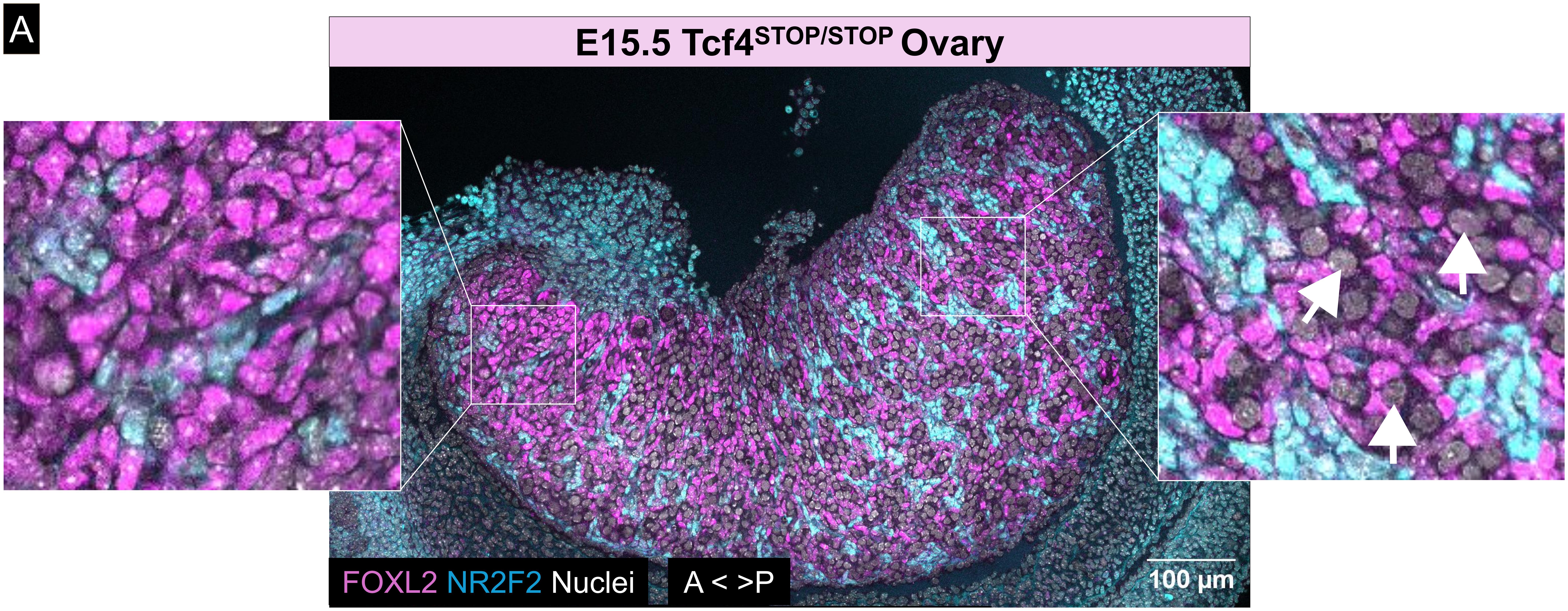


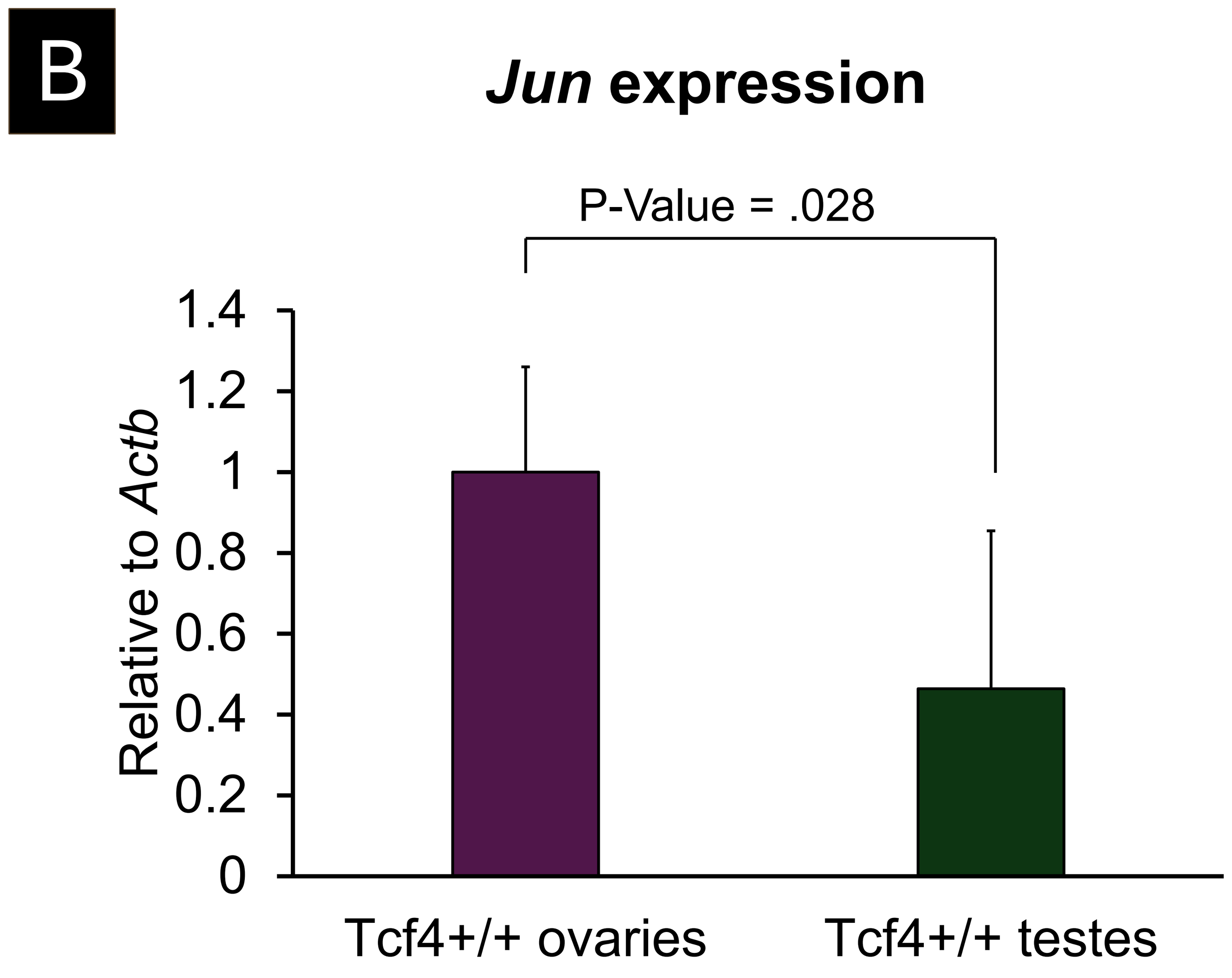
